# DrtA, a novel major facilitator superfamily transporter, contributes to intrinsic tolerance to the chemotherapeutic agent mitomycin C in *Acinetobacter baumannii*

**DOI:** 10.64898/2026.08.07.742647

**Authors:** Wuen Ee Foong, Yingqi Jin, Yumeng Duan, Haonan Su, Xuan Yan, Jiabin Huang, Heng-Keat Tam

## Abstract

Human-targeted non-antibiotic drugs are increasingly recognized for their intrinsic antibacterial activity, yet Gram-negative pathogens such as *Acinetobacter baumannii* exhibit substantial tolerance to these compounds. This tolerance is largely attributed to restricted outer membrane permeability and the activity of multidrug efflux systems. While Resistance Nodulation Division (RND) transporters have been extensively studied, the contribution of individual Major Facilitator Superfamily (MFS) transporters to non-antibiotic drug tolerance remains poorly understood. Here, we investigated H0N29_04330, designated Drug Resistance Transporter A (DrtA), a Bcr/CflA subfamily MFS transporter, to define its substrate specificity and contribution to antibiotic and non-antibiotic drug tolerance. DrtA was highly conserved across the *A. calcoaceticus-baumannii* complex and exhibited broad substrate specificity when heterologously expressed in an efflux-deficient *Escherichia coli* background, conferring resistance to benzalkonium, ethidium bromide, phenicols, and the antineoplastic agent mitomycin C. Intriguingly, *drtA* expression increased *E. coli* susceptibility to the antifolate compounds methotrexate and aminopterin, suggesting that DrtA may recognize folate-related metabolites rather than function as a dedicated antifolate transporter. In contrast, loss of *drtA* in its native *A. baumannii* host primarily impaired tolerance to mitomycin C, highlighting a context-dependent physiological role influenced by the extensive functional redundancy among *A. baumannii* efflux systems. Site-directed mutagenesis further identified M18 and the membrane-embedded protonatable residue D26 as critical determinants of DrtA transport activity and substrate recognition. Together with previous characterization of CraA, our findings demonstrate that Bcr/CflA subfamily MFS transporters contribute to protection against structurally diverse human-targeted compounds and expand the functional landscape of efflux-mediated intrinsic tolerance beyond conventional antibiotic resistance.

## 1. Introduction

Pharmaceutical agents have transformed modern medicine, yet increasing evidence indicates that many human-targeted non-antibiotic drugs possess intrinsic antibacterial activity (1. Maier et al., 2018; 2. Guillen et al., 2024). These effects are generally more pronounced in Gram-positive bacteria, whereas Gram-negative pathogens such as *Acinetobacter baumannii* display substantial intrinsic tolerance toward non-antibiotic drugs. The reduced susceptibility against non-antibiotic drugs is primarily attributed to the low permeability of the outer membrane and the efflux activity of multidrug efflux systems, particularly Resistance Nodulation cell Division (RND) transporters and Major Facilitator Superfamily (MFS) pumps, which actively extrude structurally diverse toxic compounds from the cell (3. Vila et al., 2007; 4. Laudy et al., 2016; 5. Foong et al., 2025). Beyond mediating intrinsic tolerance, these efflux pumps may also promote the emergence of antibiotic cross-resistance by enabling bacterial adaptation to non-antibiotic selective pressures (6. Ou et al., 2022). Despite growing interest in repurposing non-antibiotic drugs as antimicrobial agents, the molecular determinants governing their recognition and transport by multidrug efflux pumps remains poorly understood. In particular, the contribution of single-component MFS transporters to the extrusion of human-targeted drugs has not been systematically investigated.

Multidrug efflux pumps in *A. baumannii*, especially those belonging to the RND and MFS families, play central roles in intrinsic and acquired antibiotic resistance by exporting a wide range of antimicrobial compounds through coordinated transport networks (7. Foong et al., 2019; 8. Foong et al., 2020). While the RND systems such as AdeABC and AdeIJK are well established, the functional roles of MFS transporters remain comparatively less defined. To date, CraA is the most extensively studied MFS efflux pump in *A. baumannii* (5. Foong et al., 2025; 7. Foong et al., 2019; 9. Kröger et al., 2018), and recent evidence suggests functional interplay between MFS and RND systems, as exemplified by TetA-mediated tigecycline efflux in *A. baumannii* AYE (8. Foong et al., 2020). Consistent with the broad substrate specificity observed in the *Escherichia coli* MdfA multidrug transporter, several *A. baumannii* MFS pumps including CraA, ABAYE0913 and AmvA, display wide-ranging substrate profiles (7. Foong et al., 2019; 10. Rajamohan et al., 2010). In contrast, other MFS transporters such as AbaF, AbaQ, and CmlA5 exbihit narrower specificity, mediating resistance to fosfomycin, quinolones, and phenicols, respectively (7. Foong et al., 2019; 11. Sharma et al., 2017; 12. Pérez-Varela et al., 2018).

Previous studies have demonstrated that MFS-type transporters in *A. baumannii* possess broad and diverse substrate specificities (7. Foong et al., 2019; 5. Foong et al., 2025). Notably, the Bcr/CflA subfamily MFS transporter CraA has been shown to confer resistance to the antineoplastic agent mitomycin C (5. Foong et al., 2025). In this study, we investigate whether H0N29_04330 from *A. baumannii* ATCC19606, the homologue of ABAYE0913 from *A. baumannii* AYE, performs a similar functional role. Although previously associated with florfenicol and benzalkonium resistance (7. Foong et al., 2019), the physiological function of this transporter remains unclear. Here, we show that H0N29_04330, designated <u>D</u>rug <u>R</u>esistance <u>T</u>ransporter A (DrtA), functions as a broad-spectrum efflux pump when heterologously expressed in *E. coli*, while exhibiting a more substrate-specific role, particularly toward antineoplastic agent mitomycin C, in its native host *A. baumannii*.

## 2. Materials and Methods

### 2.1 Bacterial strains and growth conditions

All the *A. baumannii* and *Escherichia coli* strains (Supplementary Table 1) were maintained at -80°C in 20% glycerol. *A. baumannii* strains were cultured in sterile LB media while *A. baumannii* strains harbouring pBAV1K empty vector or pBAV1K-DrtA were cultured in sterile LB media supplemented with 50 mg/L kanamycin. *E. coli* S17-1 λpir donor strains harbouring the plasmid pMo130-Tel^R^_cloned-fragments were cultured in sterile LB media supplemented with 50 mg/L kanamycin. *E. coli* BW25113 Δ*emrE*Δ*mdfA* strain harbouring pET24 (empty vector) and pET24_DrtA, were cultured in sterile LB media supplemented with 50 mg/L kanamycin.

### 2.2 Cloning of MFS genes and site-directed mutagenesis

The *E. coli* codon optimized *drtA* gene with hexahistidine-tag from *A. baumannii* ATCC19606 (H0N29_04330) encoding a MFS transporter of the Bcr/CflA subfamily, was cloned into the pET24 vector using NdeI and XhoI restriction sites. The *tetA*(G) (ABAYE3637) and *craA* (ABAYE0338) genes were cloned into the pET24 vector via Gibson assembly (13. Gibson et al., 2009). Amino acid substitutions were introduced using the ExSite site-directed mutagenesis protocol (Stratagene). The integrity of the inserted gene was confirmed by DNA sequencing. All primers used for cloning are listed in Supplementary Table 2.

### 2.3 Markerless gene knockout using pMo130-Tel^R^ and conjugative transfer by biparental mating

Construction of *A. baumannii* ATCC19606 Δ*drtA* and AYE Δ*drtA* knock-out strains was performed as previously described (14. Foong et al., 2026). Briefly, approximately 1 kb DNA fragments upstream and downstream of *drtA* were amplified by PCR and assembled into suicide vector pMo130-Tel^R^ using Gibson assembly (13. Gibson et al., 2009). The resulting plasmids (Supplementary Table 1) were introduced into *E. coli* S17-1 λpir and transferred into *A. baumannii* strains by conjugation. Following conjugation, cells were incubated on LB agar at 30°C for 24 h, recovered, resuspended in 400 μl sterile LB broth, and plated onto LB agar supplemented with 30 mg/L tellurite and 50 mg/L ampicillin to select for single-crossover recombinants. Double-crossover mutants were subsequently obtained by counter-selection on LB agar containing 15% (w/v) sucrose. Correct allelic replacement was verified by PCR and confirmed by Sanger sequencing across the recombination junctions. Primer sequences are listed in Supplementary Table 2.

### 2.4 Construction of drtA complementation

For construction of the complementation plasmid pBAV1K_DrtA, PCR fragments containing the *drtA* gene and its 751 bp upstream region from ATCC19606 were cloned into pBAV1K by Gibson assembly (8. Foong et al., 2020). The resulting construct was verified by Sanger sequencing. Primers used for plasmid construction are listed in Supplementary Table 2.

### 2.5 Drug agar plate assay in E. coli and A. baumannii

Drug susceptibility assays were performed as previously described (7. Foong et al., 2019) with minor modifications. Briefly, overnight cell cultures of *E. coli* BW25113 Δ*emrE*Δ*mdfA* harbouring either the empty pET24 vector or pET24 expressing the indicated efflux pump genes, together with wildtype and Δ*drtA* strains of *A. baumannii* ATCC19606 and AYE, were adjusted to the same cell density and serially diluted (OD_600_ 10^-1^-10^-6^). Aliquots (3 μL) of each dilution were spotted onto LB agar plates supplemented with the indicated compounds (Supplementary Table 3). For *E. coli* strains, the medium was supplemented with 50 mg/L kanamycin and 0.2 mM IPTG. For complementation assays in *A. baumannii*, the medium was supplemented with 50 mg/L kanamycin. Plates were incubated overnight at 37°C before growth was assessed.

Membrane fractions of *E. coli* strains expressing DrtA and its substitution variants were prepared as previously described (15. Ma et al., 2013). Protein production was verified by Western blot analysis using a His Tag rabbit monoclonal antibody (Diagbio, China) followed by an alkaline phosphatase-conjugated goat anti-rabbit IgG (H+L) secondary antibody (APExBIO Technology, USA).

### 2.6 RNA extraction and reverse transcription qPCR

Overnight cultures of *A. baumannii* ATCC19606 were inoculated into 20 mL of LB broth to an initial OD_600_ of 0.025 and incubated at 37°C with shaking at 180 rpm until reaching an OD_600_ of 0.6-0.8. Mitomycin C was then added to a final concentration of 0.4 mg/mL, and the cultures were incubated for a further 2 h. Untreated control samples were collected immediately before mitomycin C addition. Aliquot (500 μL) of each culture were harvested and immediately stabilized with RNAstore Reagent (Tiangen Biotech Co., Ltd., China). Total RNA was extracted using the RNAprep Pure Bacteria Kit (Tiangen Biotech Co., Ltd., China).

Reverse transcription was performed using the FastKing gDNA Dispelling RT SuperMix II (Tiangen Biotech Co., Ltd., China) with 500 ng of total RNA as the template. Quantitative PCR (qPCR) was conducted using an Bio-Rad CFX Connect Real-Time PCR System (Bio-Rad Laboratories, USA) with 2× Universal SYBR Green Fast qPCR Mix (AbClonal Technology, USA) and 400 ng of cDNA per reaction. Primer sequences are listed in Supplementary Table 2, and *rpoB* was used as the reference gene. Gene expression analysis was performed as previously described (8. Foong et al., 2020). The ΔC_T_ values were calculated as C_T(rpoB)_ – C_T(targetgene)_. Differences in mean ΔC_T_ values between mitomycin C-treated and untreated control groups were assessed using an analysis of variance (ANOVA) model implemented in R v4.4.1, incorporating Gene and Gene:Treatment interaction terms. Tukey’s Honestly Significant Difference (HSD) test was subsequently applied to determine the statistical significance of differential gene expression between treatment conditions.

### 2.7 Homology models of CraA and DrtA

The CraA homology model was modeled as previously described (5. Foong et al., 2025). While the DrtA homology model was obtained via AlphaFold Protein Structure Database (16. Jumper et al., 2021). The inward-facing cavity of DrtA model was calculated using CAVER 3.0 (17. Chovancova et al., 2012).

### 2.8 Bioinformatic analysis

Homology searches for *drtA* were performed using BLASTn against *A. calcoaceticus*/*baumannii* complex (NCBI GenBank Taxonomy ID: 909768), using an E-value threshold of 0.1. The DrtA protein sequence was queried against the Comprehensive Antibiotic Resistance Database (CARD) (18. Jia et al., 2017) to identify homologous resistance determinants.

## 3. Results

### 3.1 Genomic context of H0N29_04330 across A. calcoaceticus/baumannii complex

Given the reported involvement of H0N29_04330 (homologous to ABAYE0913) in florfenicol and benzalkonium resistance, and its limited functional characterization, we first examined evolutionary conservation and genomic context of H0N29_04330. A protein homology search against the CARD (18. Jia et al., 2017) revealed that H0N29_04330 shares 31% amino acid identity with the MFS transporter Bcr-1 (Antibiotic Resistance Ontology: 3003801) (Supplementary Table 4). To determine its prevalence in *A. calcoaceticus*/*baumannii* complex, we examined a total of 38,638 genomes of *A. calcoaceticus*/*baumannii* complex available in GenBank as of March 25, 2024 (Supplementary Table 5). H0N29_04330 homologues were detected in >99.8% of genomes examined, indicating that this locus is nearly ubiquitous with the complex. The few genomes lacking H0N29_04330 may reflect incomplete or erroneous genome assemblies.

To investigate the genomic region surrounding H0N29_04330, 22 representative *Acinetobacter* genomes included in the KEGG database were selected, encompassing several widely used laboratory strains (Supplementary Table 6). Comparative genomic analysis demonstrated that the local genomic organization surrounding H0N29_04330 is highly conserved across the examined genomes (Supplementary Fig. 1). An exception was observed in *A. baumannii* ATCC17978, where a 45-kb insertion containing the fimsbactins locus (*fbsA-Q*) and multiple transposase genes separates H0N29_04330 (A1S_2584 in ATCC17978) from its upstream neighbouring gene NGG1p interacting factor NIF3, likely as a consequence of IS*5*-mediated insertion events (Supplementary Fig. 2, Supplementary Table 7-8). Across the representative genomes, *purL* encoding phosphoribosylformylglycinamidine synthase was consistently located upstream of H0N29_04330, whereas genes encoding either an NGG1p interacting factor NIF3 or a KGW-motif-containing protein were positioned downstream (Supplementary Fig. 2, Supplementary Table 9). Additional conserved genes included those encoding an acyl-CoA thioesterase, amidohydrolase, *ruvB*, *ruvA*, and a deoxyguanosine triphosphate triphosphohydrolase upstream of *purL*, as well as enoyl-CoA hydratase/isomerase and nicotinate-nicotinamide nucleotide adenylyltransferase downstream of NIF3 (Supplementary Fig. 2, Supplementary Table 9). Together, these findings demonstrate that H0N29_04330 is a highly conserved locus of the core genome of the *A. calcoaceticus/baumannii* complex.

### 3.2 Substrate profiling of DrtA and CraA in heterologous expression system

To define the substrate spectrum of H0N29_04330, the *A. baumannii* ATCC19606 gene was codon-optimized for *E. coli*, cloned into the IPTG-inducible vector pET24, and expressed in *E. coli*. The tetracycline transporter TetG and the multidrug transporter CraA were included as comparator transporters (7. Foong et al., 2019; 19. Sumyk et al., 2021). Consistent with previous reports (5. Foong et al., 2025; 7. Foong et al., 2019; 19. Sumyk et al., 2021), CraA manifested resistance to chloramphenicol and ethidium, whereas TetG mediated resistance exclusively to tetracycline (Supplementary Fig. 3A). These findings validated the use of the pET24 expression system for functional characterization of MFS transporter and established a platform for further defining the substrate profile of H0N29_04330.

In agreement with previous observations (7. Foong et al., 2019), H0N29_04330 conferred resistance to benzalkonium and florfenicol, and additionally mediated resistance to other phenicols, including chloramphenicol and thiamphenicol, as well as ethidium and mitomycin C (Fig. 1A). This observation prompted us to investigate whether H0N29_04330 could recognize additional clinically relevant non-antibiotic compounds. Unexpectedly, expression H0N29_04330 increased susceptibility to methotrexate and aminopterin, whereas neither CraA nor TetG affected susceptibility of cells to these antifolate compounds, which are widely used as chemotherapeutic and immunosuppressive agents (Supplementary Fig. 3B). To determine whether this phenotype extended to structurally related antifolate antibacterial compounds, we further examined dapsone, mafenide, sulfamethoxazole, and trimethoprim. However, H0N29_04330 did not alter susceptibility to any of these compounds (Supplementary Figure 3B), indicating that its activity toward non-antibiotic compounds is selective and primarily associated with mitomycin C and specific folate-related metabolites rather than reflecting a general antifolate transport mechanism.

**Fig. 1.**
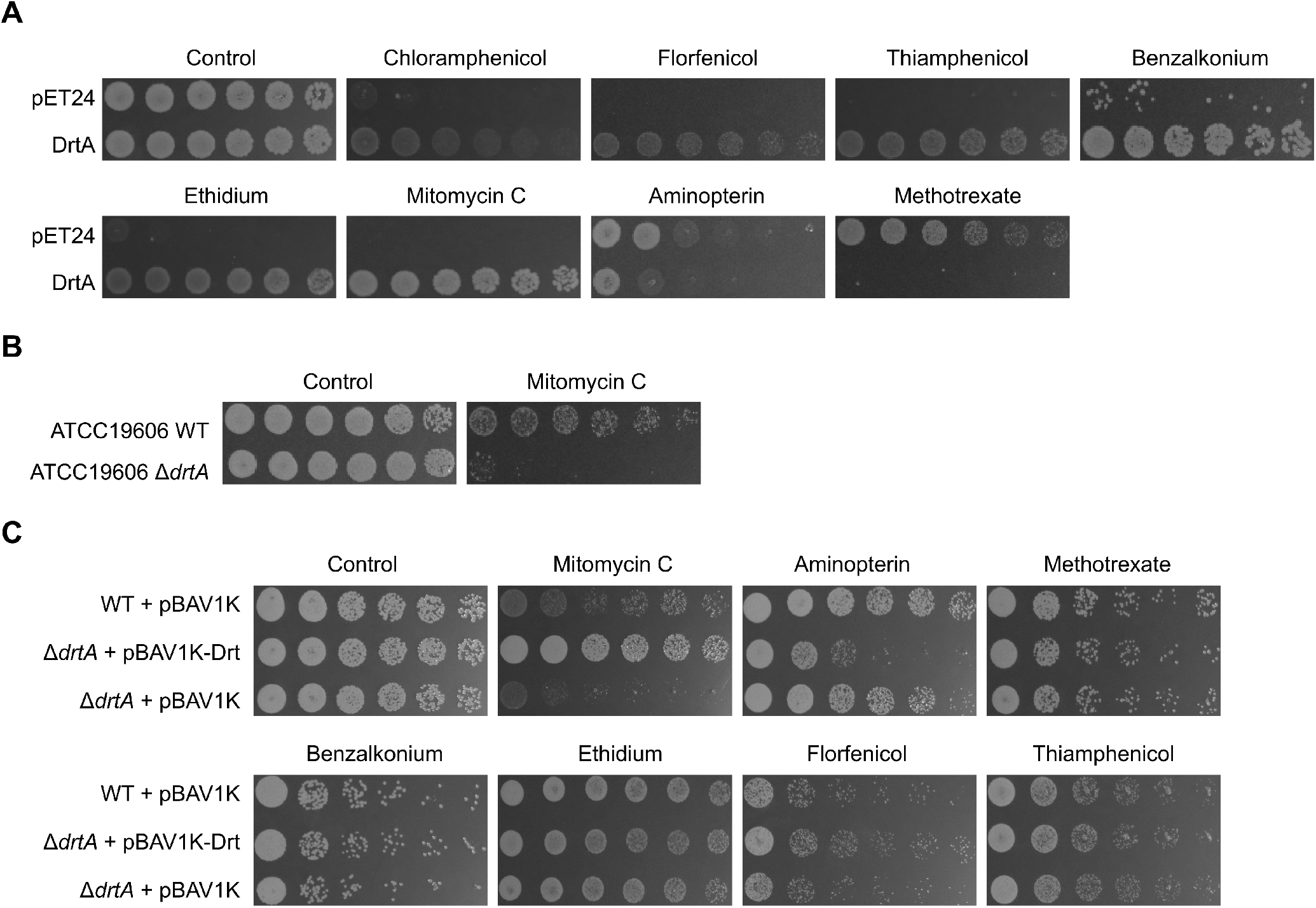
Drug resistance profiles of DrtA-overproducing *E. coli* and *A. baumannii* wildtype and Δ*drtA* strains. (A) Substrate profile of DrtA in *E. coli* BW25113 Δ*emrEΔmdfA*. (B) Mitomycin C susceptibility of wildtype and Δ*drtA* strains of *A. baumannii* ATCC19606. (C) Complementation of the Δ*drtA* mutant with *drtA* and its upstream region restores mitomycin C tolerance in *A. baumannii* ATCC19606 Δ*drtA*. Cells were serially diluted (left to right: OD_600_ 10^-1^-10^-6^) and spotted onto LB agar supplemented with the indicated compounds. Representative results from at least three independent experiments are shown.

### 3.3 Functional role of DrtA in its native host

To determine whether H0N29_04330 contributes to drug tolerance in their native hosts, a deletion mutants was generated in *A. baumannii* ATCC19606. Loss of H0N29_04330 significantly reduced tolerance to mitomycin C in *A. baumannii* ATCC19606 (Fig. 1B), whereas susceptibility to most other compounds remained unchanged (Fig. 1C). To confirm the role of H0N29_04330, a complementation construct carrying the gene together with its 751-bp putative regulatory region was introduced into the ATCC19606 Δ*drtA* mutant. Complementation restored mitomycin C tolerance and further increased susceptibility to aminopterin (Fig. 1C), confirming that both phenotypes were specifically associated with H0N29_04330 function. In the multidrug-resistant strain AYE, inactivation of the H0N29_04330 homologue, ABAYE0913, resulted in only a modest increase in mitomycin C susceptibility (Supplementary Figure 4) Together with previous studies of CraA (5. Foong et al., 2025; 7. Foong et al., 2019), our findings indicate that MFS transporter in *A. baumannii* possesses broader and more diverse substrate repertoires than previously recognized. Accordingly, H0N29_04330 was designated <u>D</u>rug <u>R</u>esistance <u>T</u>ransporter A (DrtA).

To determine whether *drtA* expression is responsive to mitomycin C in *A. baumannii*, we quantified the transcript levels of *drtA* by RT-qPCR, using *craA* as a positive control because CraA plays a key role in mitomycin C tolerance (5. Foong et al., 2025). As anticipated, *craA* expression was significantly upregulated following mitomycin C exposure (ΔΔC_T_ = 4.32 ± 0.26) (Supplementary Figure 5). In contrast, *drtA* was expressed at a low basal level (ΔC_T_ = - 6.13 ± 0.18) and showed minimal induction in response to mitomycin C (ΔΔC_T_ = 0.37 ± 0.26). These findings suggest that the low basal expression and limited inducibility of *drtA* may explain why deletion of *drtA* had little effect on susceptibility to most compounds tested. In contrast, the strong induction of *craA* in response to drug exposure may compensate for the loss of *drtA* (5. Foong et al., 2025; 14. Foong et al., 2026), thereby masking the phenotypic consequences of the *drtA* knockout.

### 3.4 Functional analysis of membrane-embedded residues

Sequence alignment of DrtA revealed that, unlike typical MdfA subfamily MFS transporters, it contains a single membrane-embedded proton-titratable residue, D26, which is conserved in CraA, MdfA, and the chloramphenicol transporter CmlA (Fig. 2A) (7. Foong et al., 2019). In addition, DrtA lacks the second conserved proton-titratable residue in CraA (E38) and MdfA (E26), where this position is replaced by M18. Although primary sequence alignment suggested N328 as a potential counterpart of CraA E338, structural alignment of DrtA and CraA identified F322 as the most likely equivalent residue (Fig. 2B). Therefore, F322 was included in the mutational analysis. In addition, residues S22 and I25, corresponding to Y42 and N45 in CraA, respectively, were selected based on their proposed involvement in substrate recognition in CraA. To assess the functional importance of these residues, D26, M18, and F322 were substituted to alanine or glutamate, while S22 and I25 were substituted with alanine. The resulting DrtA variants were evaluated using drug susceptibility assays in *E. coli* BW25113 Δ*emrE*Δ*mdfA* expressing either wildtype DrtA or the corresponding substitution variants (Fig. 3). All *drtA* mutants were expressed equally well compared with wildtype *drtA* from the same expression vector (Supplementary Figure 6).

**Fig. 2.**
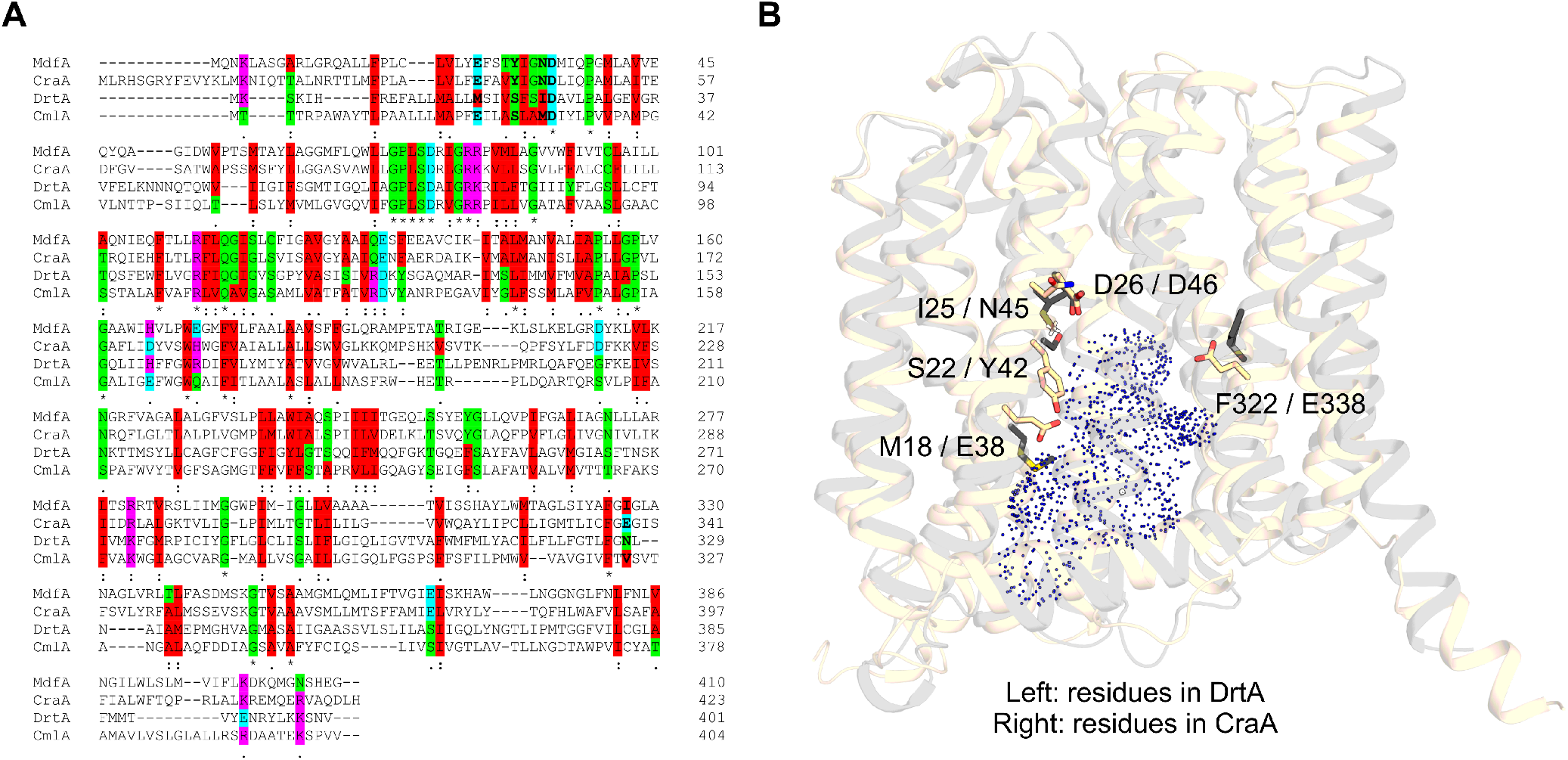
Protein sequence alignment and structural comparison of CraA, DrtA, and related MFS transporters. (a) Multiple sequence alignment of CraA, MdfA, DrtA, and CmlA. MdfA, *Escherichia coli* MdfA; CraA, *A. baumannii* AYE CraA (GenBank accession number: CAJ77876); CmlA, *A. baumannii* AYE CmlA (GenBank accession number: CAM88409); DrtA, *A. baumannii* AYE DrtA (GenBank accession number: CAM85858). Residues selected for mutational analysis are highlighted in bold. Amino acid residues are colour-coded according to their physicochemical properties: hydrophobic residues, red; negatively charged residues, cyan; hydrophilic residues, proline, cysteine, and glycine, green; positively charged residues, magenta. (b) Structural superimposition of DrtA and CraA. The AlphaFold-predicted structure of DrtA and the CraA homology generated based on the MdfA_Q131R_L339E variant (PDB: 6EUQ), as described previously (5. Foong et al., 2025), are shown in cartoon representation. CraA and DrtA are coloured beige and grey, respectively. Residues selected for amino acid substitution are shown as sticks and coloured according to their corresponding protein model. The inward-facing cavity was calculated using CAVER 3.0 (22. Chovancova et al., 2012) and is shown as a dot-surface representation.

**Fig. 3.**
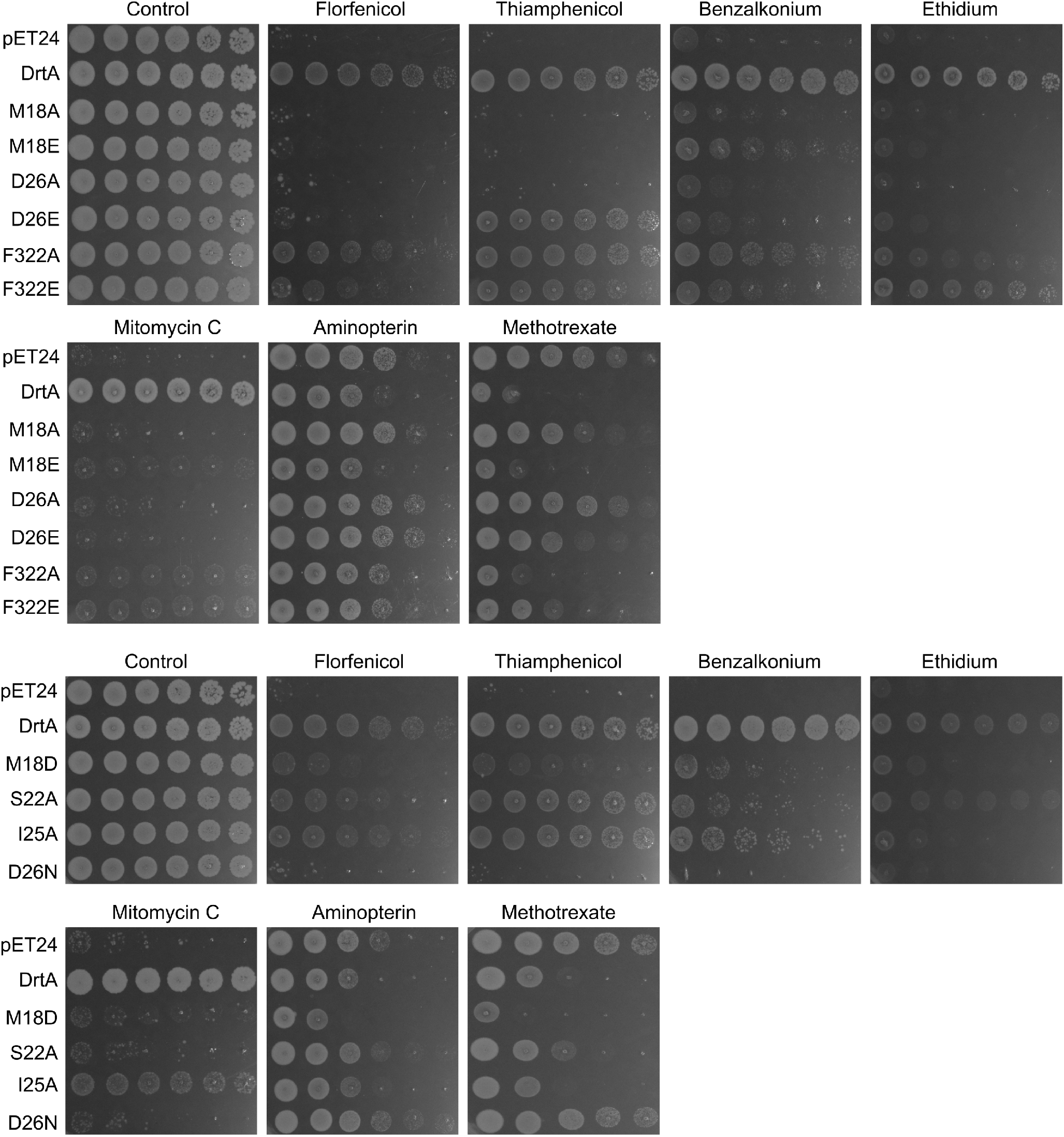
Drug resistance profiles of *E. coli* BW25113 Δ*emrE*Δ*mdfA* harboring DrtA variants with mutation of selected residues lining along the predicted binding pocket. Cells were serially diluted (left to right: OD_600_ 10^-1^-10^-6^) and spotted onto LB agar supplemented with the indicated compounds. Representative results from at least three independent experiments are shown.

As anticipated, substitution of D26 with either alanine, glutamate or the protonated mimic asparagine abolished DrtA-mediated resistance to most tested drugs, with the exception that the D26E variant restored resistance to thiamphenicol. These findings suggest that the protonatable carboxylate residue at this position is critical for transport activity, and that extension of the side chain by one methylene group can partially preserve transport function in a substrate-dependent manner. Similarly, substitution of the M18 with alanine abolished resistance to all tested compounds, whereas the M18E variant retained the wildtype susceptibility profile toward methotrexate. In contrast, substitution at S22, I25, and F322 were generally tolerated, indicating that this residue is not essential for the transport activity of DrtA under the conditions tested. Together these results suggest that D26 plays a central role in proton coupling and energy transduction during DrtA-mediated transport, whereas M18 may contribute to substrate recognition or binding. The differential effects of these mutations compared to CraA and MdfA further highlight the importance of individual residues and in determining substrate-specific transport activity.

## 4. Discussion

Our findings unveil H0N29_04330, hereafter designated <u>D</u>rug <u>R</u>esistance <u>T</u>ransporter A (DrtA), as an MFS efflux pump with a prominent role in mediating tolerance to the antineoplastic agent mitomycin C in its native host (Fig. 1C, Supplementary Fig. 4). Although DrtA was previously associated primarily with florfenicol and benzalkonium tolerance (7. Foong et al., 2019), our results substantially expand its substrate profile. When heterologously expressed in *E. coli*, DrtA conferred resistance to additional phenicols, ethidium, and the human-targeted compounds mitomycin C, while also modulating susceptibility to methotrexate, and aminopterin (Fig. 1), thereby extending its functional relevance beyond conventional antimicrobial compounds. Importantly, the broad substrate profile observed in *E. coli* contrasted with the more restricted physiological role of DrtA in *A. baumannii*, where its contribution was primarily limited to mitomycin C tolerance (Fig. 1C, Supplementary Fig. 4). This species-dependent difference is likely attributable to the low basal expression and limited inducibility of *drtA*, together with the extensive functional redundancy among efflux systems in *A. baumannii*. In *A. baumannii*, RND and MFS transporters, including AdeABC, AdeIJK, AmvA, and CraA, collectively mediate tolerance to structurally diverse compounds such as benzalkonium, phenicols, ethidium, and mitomycin C (7. Foong et al., 2019; 10. Rajamohan et al., 2010; 20. Magnet 2001; 21. Damier-Piolle 2008). Consequently, loss of a single transporter may have only a modest phenotypic impact under standard laboratory conditions. By contrast, *E. coli* possesses a comparatively limited multidrug efflux repertoire, consisting primarily of AcrAB-TolC, EmrE, and MdfA, which likely accentuates the substrate profile of DrtA during heterologous expression (22. Alon Cudkowicz et al., 2019).

One intriguing observation was that overexpression of *drtA* rendered *E. coli* highly susceptible to methotrexate and aminopterin, but not other structurally related antifolate compounds (Fig. 1B, Supplementary Fig. 3B). Similarly, complementation of *drtA* in *A. baumannii* using the high-copy-number pBAV1K plasmid increased susceptibility to aminopterin (Fig. 1C) (23. Bryksin et al. 2010), whereas the Δ*drtA* mutant showed no significant change in aminopterin susceptibility. These findings suggest that DrtA overproduction, rather than the physiological level of *drtA* expression, contributes to increased aminopterin sensitivity. Interestingly, substitution of the protonatable membrane-embedded residue D26 or the M18 (only M18A) in DrtA abolished methotrexate-associated susceptibility, while simultaneously eliminating transport activity toward other tested substrates. These findings raise the possibility that DrtA may facilitate the uptake or intracellular accumulation of methotrexate and aminopterin, rather than simply increasing membrane permeability as a consequence of transporter overproduction. Nevertheless, the biological significance of this potential transport activity remains unclear. Given the structural similarity of these compounds to folate (Supplementary Table 3), we speculate that DrtA may recognize folate or folate-derived metabolites rather than function as a dedicated antifolate transporter. This hypothesis is supported by previous evidence that the *E. coli* AbgT transporter mediates uptake of the folate breakdown product para-aminobenzoate-glutamate (24. Maynard et al. 2018), suggesting that multidrug MFS transporters may contribute to the trafficking of folate-related metabolites in addition to xenobiotic compounds.

Collectively, these findings reinforce the concept that MFS transporters contribute not only to antibiotic resistance but also to the intrinsic tolerance of *A. baumannii* toward structurally diverse human-targeted drugs. Together with the RND efflux systems AdeABC, AdeIJK, and AdeFGH (7. Foong et al., 2019; 8. Foong et al., 2020), DrtA and CraA likely constitute an integrated efflux network that protects *A. baumannii* against both antibiotic and non-antibiotic toxicants. Such functional overlap may contribute to the remarkable adaptability of this pathogen and may provide a mechanistic basis for the emergence of cross-resistance between clinically used non-antibiotic drugs and antimicrobial agents. Further investigation of efflux-mediated non-antibiotic tolerance will improve our understanding of intrinsic resistance mechanisms and may uncover new opportunities for therapeutic intervention.

## CRediT authorship contribution statement

**Wuen Ee Foong:** Writing – review & editing, Writing – original draft, Supervision, Investigation, Methodology, Visualization, Validation, Data curation, Funding acquisition, Conceptualization. **Yingqi Jin:** Investigation. **Yumeng Duan:** Investigation. **Haonan Su:** Investigation. **Xuan Yan:** Investigation. **Jiabin Huang:** Writing – review & editing, Investigation, Methodology, Visualization, Validation. **Heng-Keat Tam:** Writing – review & editing, Writing – original draft, Supervision, Project administration, Data curation, Visualization, Supervision, Investigation, Methodology, Funding acquisition, Conceptualization.

## Acknowledgements

This work was supported by grants from the start-up package through the University of South China (221RGC012), the Hunan Provincial Natural Science Fund (2024JJ5326), the Science and Technology Innovation Program of Hunan Province (2024RC9011), and the National Natural Science Foundation of China (W2532023).

## Supporting information

Supplementary data associated with this article can be found in Supplementary Information.

