## Supplementary Figure 1 to Supplementary Figure 6 for "DrtA, a novel major facilitator superfamily transporter, contributes to intrinsic tolerance to the chemotherapeutic agent mitomycin C in *Acinetobacter baumannii*"

**Supplementary Figure 1.** The *drtA* (ABAYE0913 in *A. baumannii* AYE and H0N29_04330 in *A. baumannii* ATCC19606) loci in *A. baumannii* isolates. The sequence identity between genes is shown by shading according to the identity scale bar.

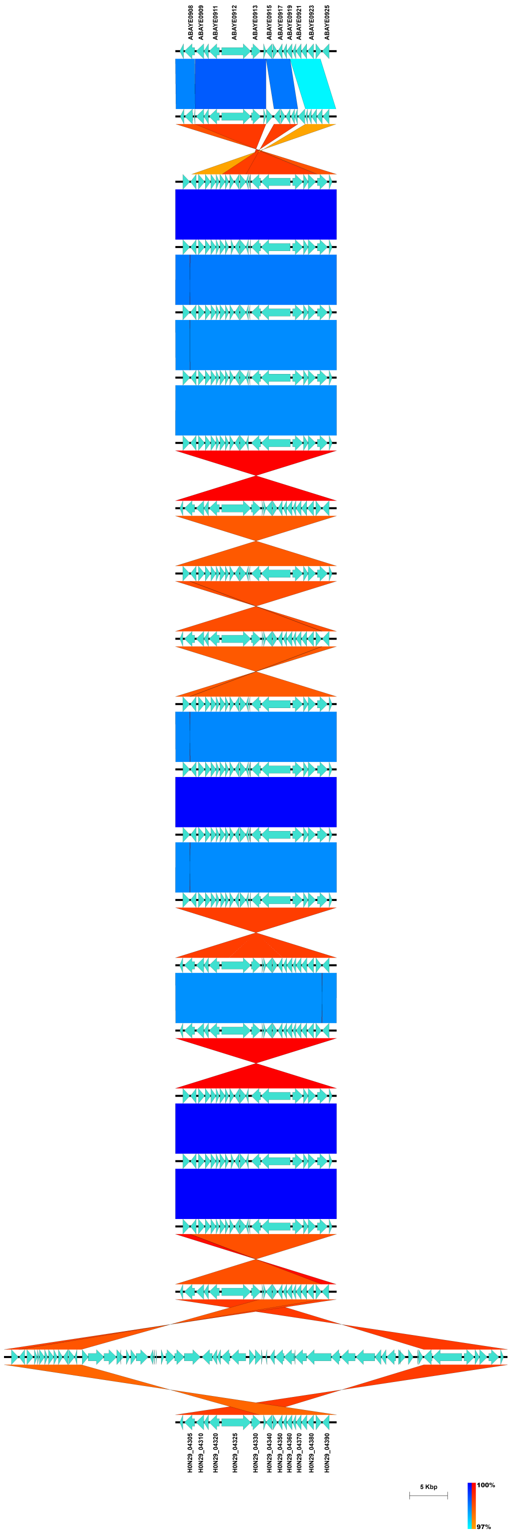

**Supplementary Figure 2.** Insertion of 45-kbp fragment into *drtA* genomic locus of *A. baumannii* ATCC17978. Genes are represented by arrows coloured according to similarity groups, with grey arrows indicating genes not part of any similarity group. The sequence identity between genes in the same similarity group is indicated by shading, according to the identity scale bar. Detailed gene annotations are shown and colour-coded.

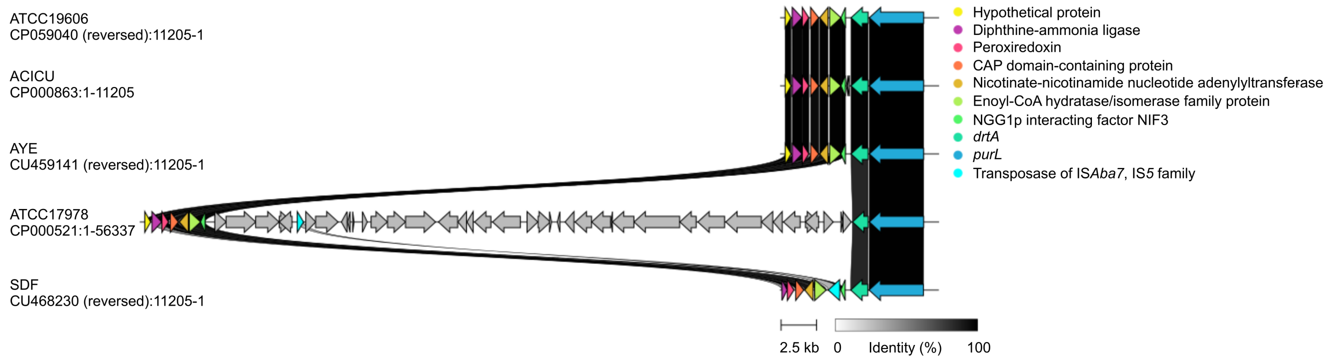

**Supplementary Figure 3.** Drug resistance profiles of MFS transporter-overproducing *E. coli* strains. (A) Drug susceptibility profiles of CraA-, TetG-, and DrtA-overproducing *E. coli* BW25113 Δ*emrE*Δ*mdfA* harbouring pET24 (empty vector) or pET24 expressing the corresponding efflux pump genes. (B) Antifolate susceptibility profile of DrtA in *E. coli* BW25113 Δ*emrE*Δ*mdfA*. Cells were serially diluted (left to right: $OD_{600}$ $10^{-1}$-$10^{-6}$) and spotted onto LB agar supplemented with the indicated compounds. Representative results from at least three independent experiments are shown.

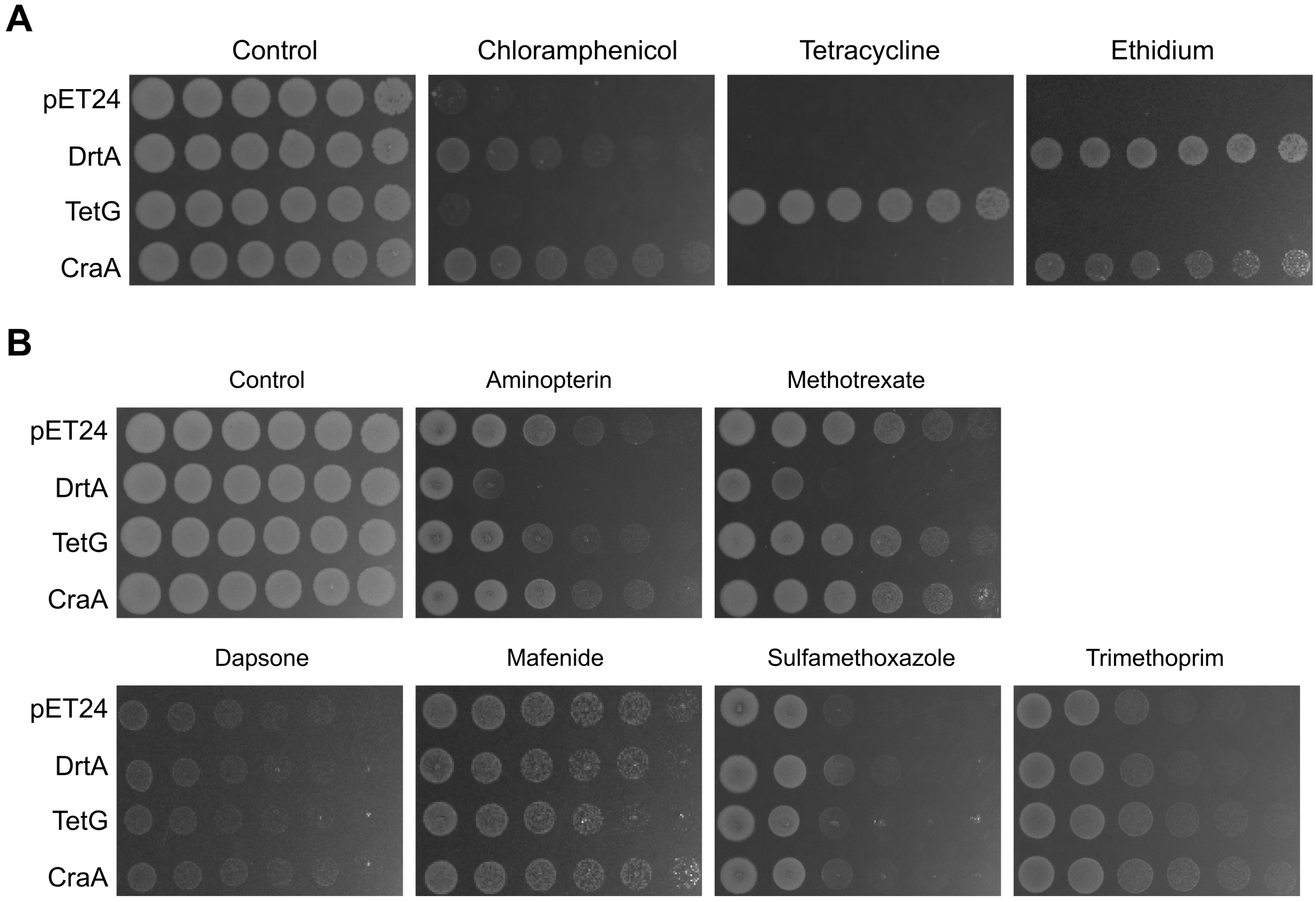

**Supplementary Figure 4.** Mitomycin C susceptibility of wildtype and Δ*drtA* strains of *A. baumannii* ATCC19606. Representative results from at least three independent experiments are shown.

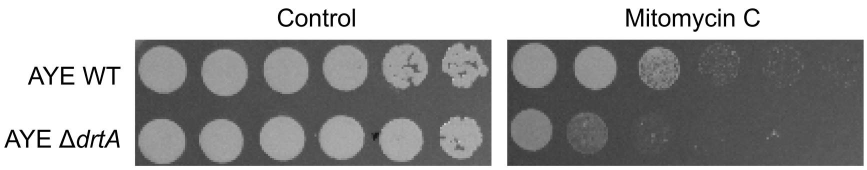

**Supplementary Figure 5.** Gene expression analysis of *drtA* gene in *A. baumannii* ATCC19606 in the absence or presence of mitomycin C (MIT). The untreated sample (without mitomycin C) was used as the calibrator for $\Delta\Delta C_T$ analysis. Data points and error bars represent the mean ± SEM from ≥ 5 independent biological replicates. Mitomycin C did not significantly induce *drtA* expression ($\Delta\Delta C_T$ = 0.37 ± 0.26, fold change = 1.3), but induce *craA* expression ($\Delta\Delta C_T$ = 4.32 ± 0.26, fold change = 20.0, and Tukey's Honest Significant Differences test, $p < 0.001$).

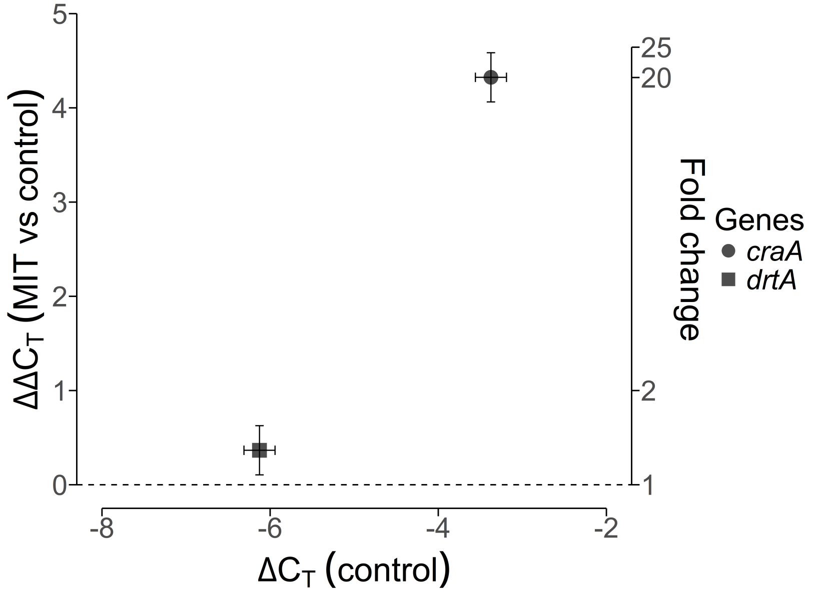

**Supplementary Figure 6.** The Western blot analysis of the whole cell extracts of wildtype DrtA and its variants confirms that all DrtA variants expressed equally well compared to wildtype CraA.

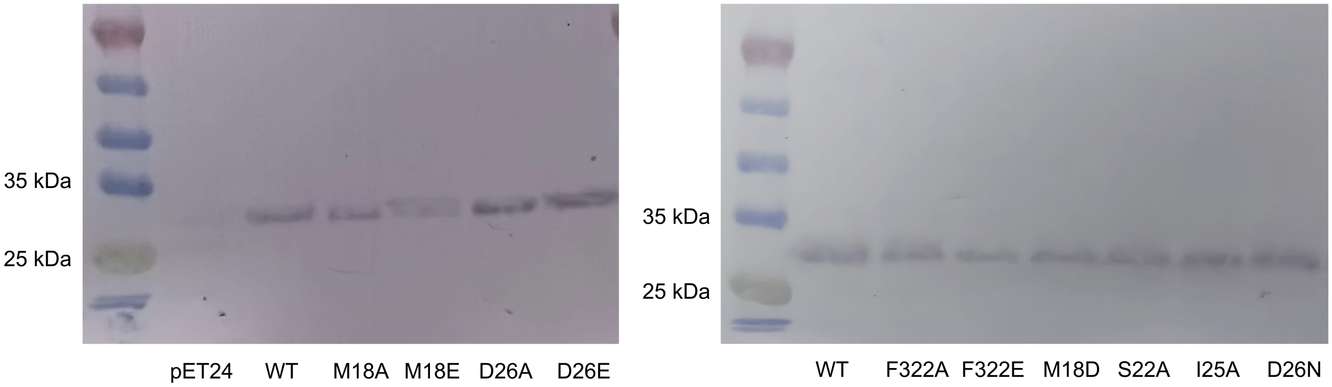
